# Identification of pyroptosis-related diagnostic model relate to immune infiltration in osteoarthritis

**DOI:** 10.1101/2025.02.13.638031

**Authors:** Meimei Xiao, Yue Qiu, Jinzhi Meng, Cancai Jiang, Yanming Chen, Jun Yao

## Abstract

**Background:** Osteoarthritis (OA) is a common chronic diseases which were associated with aging and progressive joint dysfunction. Pyroptosis, a novel mode of cell death. The effect of pyroptosis on osteoarthritis (OA) progression is still in the exploratory stage, and the specific mechanism is unclear.

**Methods:** OA data sets were obtained from the GEO databases website. Bioinformatics analysis was conducted to identify pyroptosis-related genes (PRGs), construct a pyroptosis-related diagnostic model, and identify subtypes. We established the random forest (RF) model as well as a nomogram of differentially expressed PRGs. The association between immunity and pyroptosis was comprehensively explored. Finally, the expression levels of 3 OA characteristic genes were verified by qRT-PCR and immunohistochemistry.

**Results:** Seven differentially expressed PRGs (CASP3, GPX4, NOD1, TIRAP, SCAF11, CASP4, and CASP9) were obtained by differential analysis to establish RF models, and three important OA genes (CASP9, TIRAP, and SCAF11) were obtained. ROC curve proved that the established model had high diagnostic accuracy. Based on the above three important PRGs, a high-precision nomogram model is constructed. Two PRG models are determined by consensus clustering method. The enrichment degree of immune cells in cluster A was higher than that in cluster B, suggesting that cluster A might be related to the occurrence and development of OA. GPX4 was found to be positively correlated with higher immune infiltration. Finally, the expression levels of 3 OA characteristic genes were verified by qRT-PCR and immunohistochemistry.

**Conclusion:** We constructed an effective prognostic model according to PRGs in OA and integrated explored the association between pyroptosis and immunity-related elements. This article provides a strong point for investigating directed immunotherapy methods with novelty for OA.

## 1 Introduction

Osteoarthritis (OA) is a persistent inflammatory joint disorder characterized by cartilage degradation, synovial inflammation, as well as subchondral bone remodeling, which limits the movement of middle-aged and older adults (1, 2). Many factors, such as heredity, environment, metabolism and mechanical stress, and complex pathogenesis, may influence osteoarthritis. It is estimated that among middle-aged and older adults over the age of 60, the probability of men suffering from knee OA is about 10% and that of women is about 13%, which makes knee osteoarthritis become the main cause of physical disability in middle-aged and older adults(3). Because the pathological mechanism of OA is still being determined, there is still no breakthrough in the early treatment of osteoarthritis. Finally, only joint replacement surgery can be performed. Therefore, exploring the exact pathological mechanism of osteoarthritis has become an urgent problem in the early treatment of OA.

Apoptosis and autophagy are two major mechanisms in the pathogenesis of osteoarthritis. At the same time, pyroptosis is a kind of programmed cell death involved in inflammatory that differs from them and has been confirmed to be related to the pathological development of OA (4, 5). Recent studies have found that pyroptosis, a form of programmed inflammatory cell death, may play an essential role in this chronic inflammatory disease. The role of synovial and chondrocyte pyroptosis in the development of osteoarthritis has been reported. NF-κB, P2X7, Nrf2 and HIF-1α may play an essential role in the pyroptosis of OA cells mediated by NLRP inflammatome (6, 7). The research on the role of pyroptosis in the pathology of OA has gradually become a new hot topic. The typical characteristic of pyroptosis is that it is closely dependent on the activation of some caspase, accompanied by the secretion of pro-inflammatory cytokines as well as the cleavage of gasdermin D (GSDMD), leading to inflammation (8). Pyroptosis differs from other forms of cell death, especially in morphological and biochemical properties, such as the characteristics of apoptotic cells. However, the study of OA-related pyroptosis is still preliminary, and its specific regulatory mechanism is unclear.

Therefore, we aim to investigate the feasible role of pyroptosis in OA as well as provide the choice of path theoretically for exploring new clinical therapies for OA.

## 2 Material And Methods

### 2.1 Data processing and identification of pyroptosis-related genes

The gene expression profiles of GSE82107, GSE10575, GSE16464, GSE27390 and GSE19060 were obtained from Gene Expression Omnibus (GEO) (https://www.ncbi.nlm.nih.gov/geo/), including 71 samples from patients with OA (n=46) and normal control (n=25). To identify potential pyroptosis-related genes (PRGs) between normal and OA samples, we used the R “limma” package to extract differentially expressed PRGs. PRGs with logFC absolute value > 1 as well as P < 0.05 were regarded as significant differentially expressed PRGs. The position of PRGs on chromosomes was depicted. Also, the relationship between PRGs was displayed.

### 2.2 Construction of the models

The random forest (RF) and support vector machine (SVM) models were established according to the obtained PRGs. We utilized the RF and SVM models to indicate the incidence of OA and offer accurate results. Also, we used the reverse cumulative distribution of residuals, boxplots, as well as ROC curves to evaluate the prediction model’s degree of preciseness and accuracy. We utilized the “RandomForest” package in R software to construct the RF model and identify the significant PRGs to expect the incidence of OA. The values of ntree and mtry in the RF model were set to 100 and 3, respectively. Subsequently, the candidate PRGs were selected with the threshold of 10-fold cross-validation.

### 2.3 Analysis of the risk score model and construction of the nomogram

The “rms” packages constructed a nomogram to determine the disease risk for OA patients. The nomogram was constructed for TIRAP, CASP9, and SCAF11. We evaluate the model’s accuracy and prediction value via the decision curve analysis (DCA) as well as clinical impact curves. Also, we established the calibration curves to display the distinction of the predicted as well as real effects of the nomogram.

### 2.4 Establish the PRG signature via principal component analysis (PCA)

PCA was performed via the “Consensus ClusterPlus” package to explore the stratification of OA patients with diverse PRG molecular subtypes.

### 2.5 The exploration of immune cells infiltration

The arithmetic of single-sample gene set enrichment analysis (ssGSEA) is a method commonly used to calculate immune cell infiltration. This method estimates the relative enrichment of each gene set in that sample by comparing the gene expression data of each sample with a specific gene set (the immune cell gene set). In our research, the correlation between different PRGs and immune cell enrichment was explored via immune correlation investigation. PRGs significantly associated with immune cell infiltration were extracted for subsequent analysis. The above-extracted PRGs were subgrouped into groups with high and low expression according to the median, respectively. The histogram displayed the immune cell infiltration of the high as well as low-expression groups.

### 2.6 Identification of different PRG patterns as well as analysis of enrichment function

The “limma” package in R software was utilized to select DEGs of different PRG patterns based on the standard q value filter < 0.05. The Gene Ontology (GO), as well as Kyoto Encyclopedia of Genes and Genomes (KEGG) enrichment analysis of PRGs, were conducted via the “clusterProfiler” package in R software, and items with p-value< 0.05 as well as false discovery rate (FDR) < 0.05 were regarded as statistically significantly items.

### 2.7 Culture and extraction of rat chondrocytes

We aseptically obtained the knee cartilage of SD rats aged 3-7 days and then isolated and cultured chondrocytes in vitro. The mixture was divided into small pieces and digested with trypsin (Solabio, China) at 37°C for 30min. Then we combined it with collagenase II (1 mg/ml; Gibco) and high glucose DMEM (Gibco, USA) to incubate for 6 hours.And chondrocytes were exclusively collected after centrifugation at the speed of 1500rpm/3min. Gibco (Gibco, USA) and penicillin/streptomycin (Solarbio, China) were used in the DMEM culture medium. Rat chondrocytes were treated with lipopolysaccharide (LPS) to establish an OA cell model. We got approval from the Medical Ethics Committee of the First Affiliated Hospital of Guangxi Medical University (Approval Number: 2023-E585-01).

### 2.8 RNA extraction and qRT-PCR

We extracted total RNA in cells using Hipure Total RNA Minikit (Magen, China), following the manufacturer’s instructions. PrimeScript RT kit and gDNA eraser (Takara, China) were used to reverse-transcribe the extracted RNA into cDNA. Quantitative reverse transcription-polymerase chain reaction (qRT-PCR) was carried out using LightCycler 480(Roche, Germany) and Fast Start Universal Sybr Green Master Mix (Roche, Germany) under the following circumstances: 10 minutes at 95°C, 15 seconds at 95°C and 1 minute at 60°C. Data from dissolution curves were utilised to confirm the specificity of PCR. We obtained the relative expression levels of genes (CASP9, SCAF11 and TIRAP) by the 2-DDCt method. The target mRNA level was standardized to glyceraldehyde-3-phosphate dehydrogenase level (GAPDH). Then they were compared with that of the control group.

### 2.9 Immunohistochemistry

The expressions of CASP9, TIRAP and SCAF11 were detected by immunohistochemical staining. The donor’s consent to use human cartilage specimen in this study was approved by the Medical Ethics Committee of the First Affiliated Hospital of Guangxi Medical University (Approval number: 2023-E585-01).

After washing human cartilage samples in PBS, the cells and sections were exposed to 3%(v/v) hydrogen peroxide at room temperature for 15 minutes to block any endogenous peroxidase activity. The cells and areas were then blocked with 10% normal goat serum at room temperature for 20min and interacted with CASP9, TIRAP, and SCAF11 primary antibodies. The samples were then cultured with secondary antibodies and biotin-labeled horseradish peroxidase. Before hematoxylin restaining, antibody binding was observed with 3,3 ’-diaminobenzidine tetramine hydrochloride kit (ZSBG-BIO, China). After gradual dehydration and sealing with a neutral resin, the cells were observed and captured with an inverted phase contrast microscope (Olympus BX53).

### 3.0 Statistical analysis

As mentioned, we conducted statistical analyses via Perl software and R software (version 4.1.3). Unless otherwise specified, items with a p-value < 0.05 were generally regarded as statistically significant items.

## 3 Results

### 3.1 The landscape of the 7 PRGs in OA

Seven differentially expressed PRGs were obtained (Figures 1A, B). CASP3, GPX4, NOD1, TIRAP, SCAF11, CASP4 and CASP9 expressions were significantly lower in OA patients. After data visualization of candidate PRGs’ position in the chromosome, figure 1C displayed the corresponding positions in the chromosome of the obtained seven PRGs.

**FIGURE 1.**
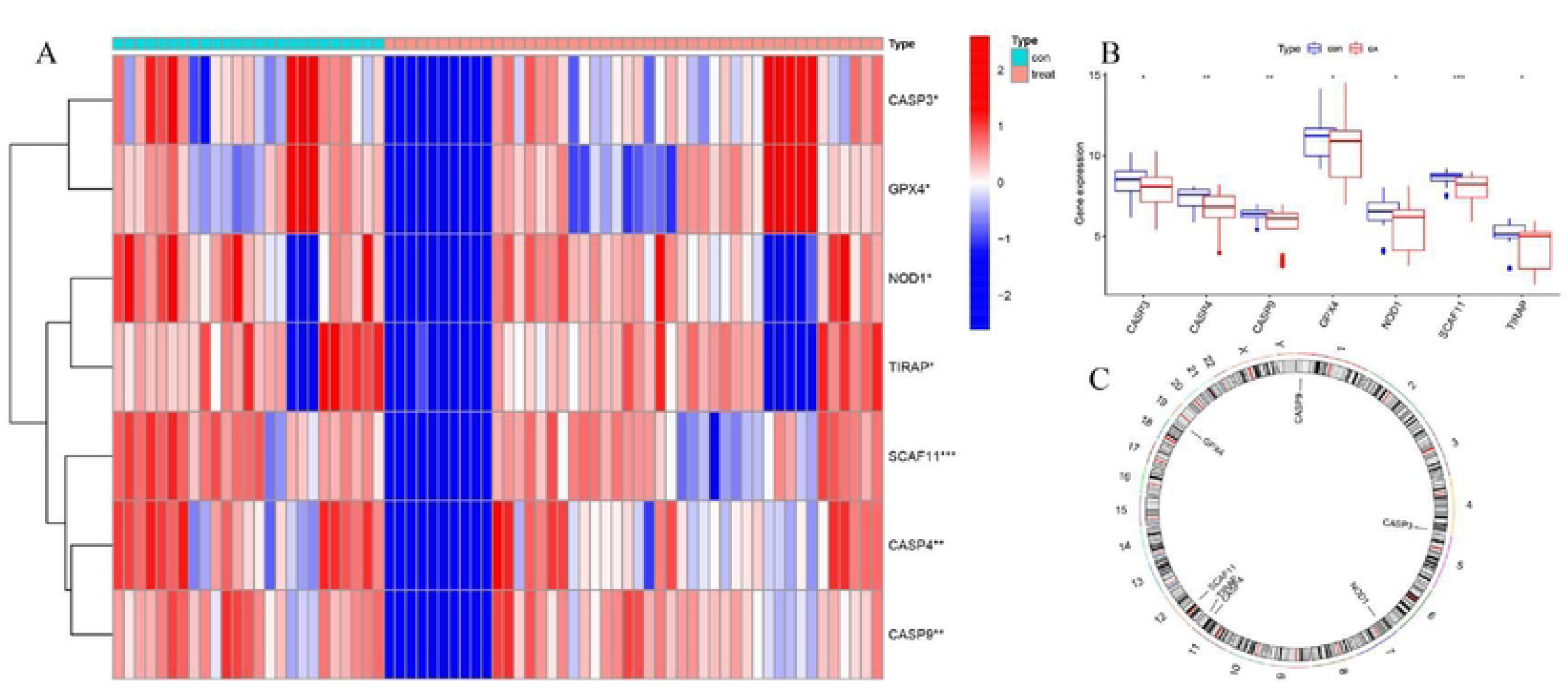
General situation of differential expression of PRGs between normal samples and OA patients. (A) 7 PRGs expression heat maps between the two groups; (B) 7 PRGs expression histograms between the two groups; (C) The chromosomal locations of 7 PRGs. *P<0.05; **P<0.01; ***P<0.001.

### 3.2 The relationship between PRGs

To elucidate the association between differentially expressed PRGs in OA patients, we conducted a linear regression analysis via R software (Figure 2A–G). The results indicated that NOD1 was positively correlated with CASP4 and CASP9. SCAF11 was positively correlated with CASP3, CASP4 and CASP9. TIRAP was positively correlated with CASP4 and CASP9.

**FIGURE 2.**
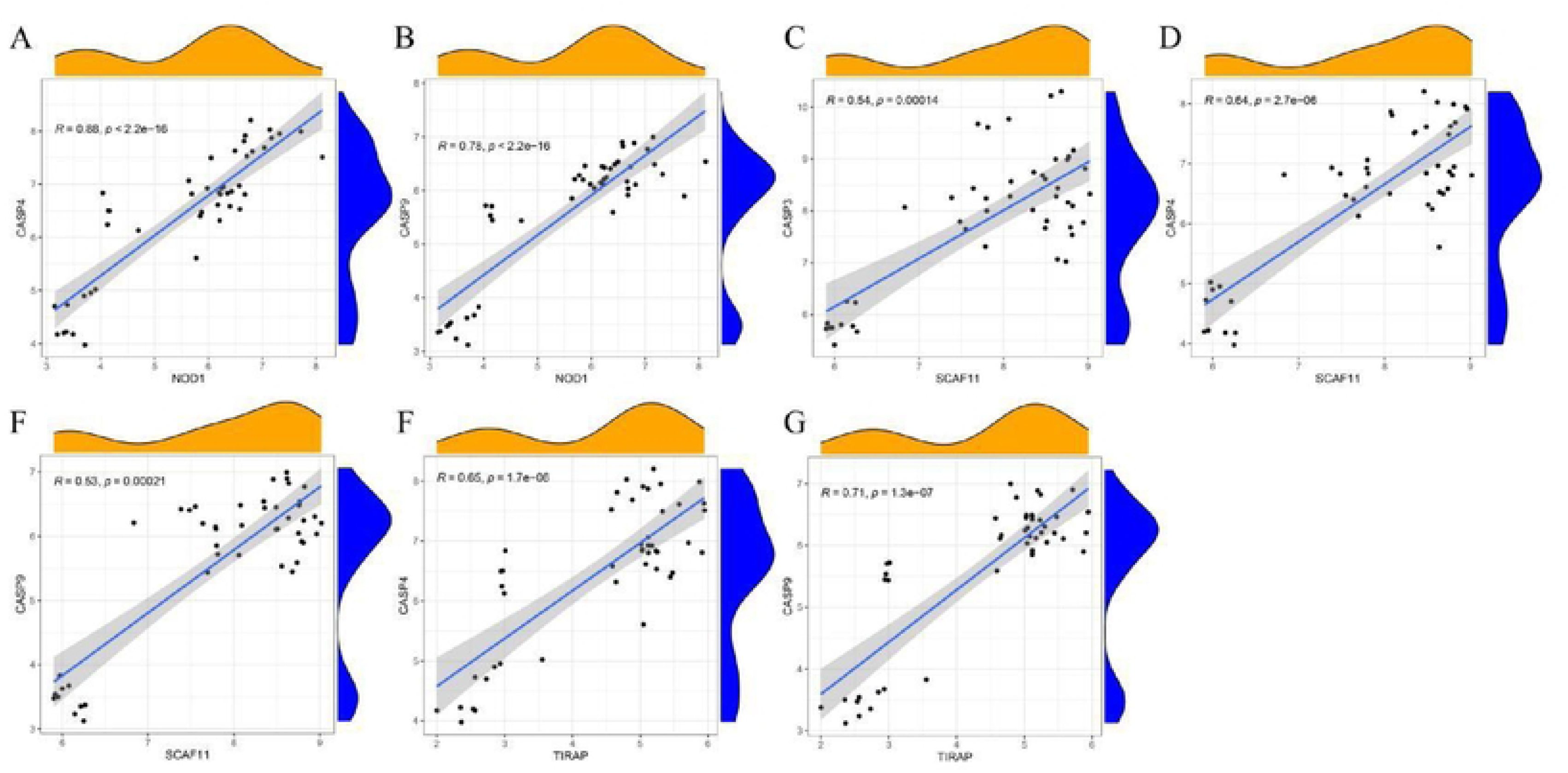
Correlation between PRGs in OA. (A) NOD1 and CASP4; (B) NOD1 and CASP9; (C) SCAF11 and CASP3; (D) SCAF11 and CASP4; (E) SCAF11 and CASP9; (F) TIRAP and CASP4; (G) TIRAP and CASP9.

### 3.3 Establishment of the RF and SVM models

The RF and SVM models were established according to the above seven differential genes to describe OA characteristics. The results of the residual boxplot of the two models (Figure 3A) as well as the line chart of the reverse cumulative distribution of residuals (Figure 3B) of the two models were consistent; both of them indicated the RF model was of less residuals, which favored the RF model as the selected one compared with the SVM model. Simultaneously, the area under the curve (AUC) in the ROC curve also favored the RF model compared with the SVM model (Figure 3C). Also, the error values were shown on the cross-validation curve with 10-fold of the treatment, control groups as well as the overall sample (Figure 3D). Also, Figure 3E visualizes the above 7 PRGs on the random decision forest map, and the results suggest that TIRAP, SCAF11 and CASP9 greatly influence this model and can be considered an important indicator for constructing a nomogram. We concluded that the RF model was a considerable model to expect the occurrence of OA.

**FIGURE 3.**
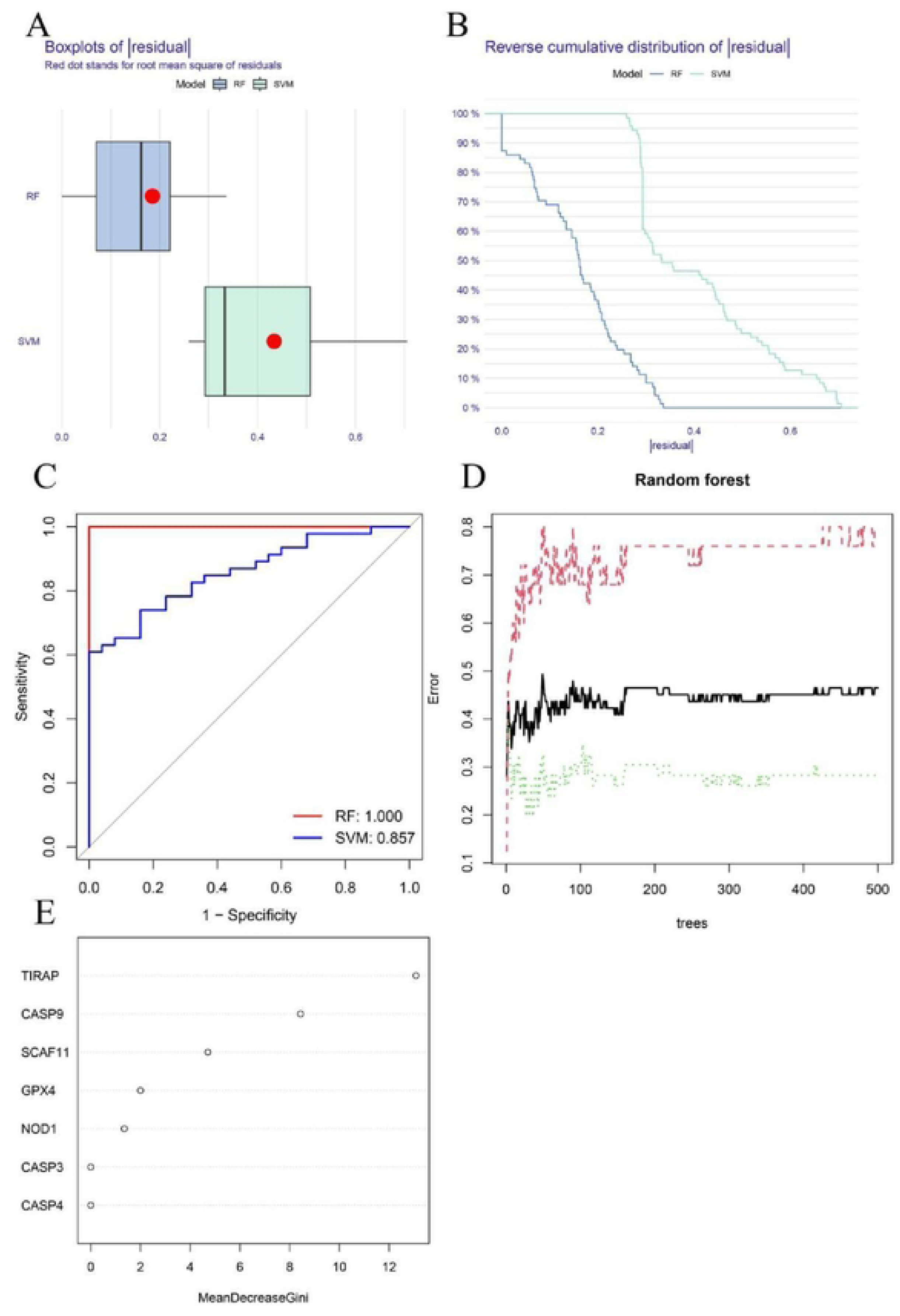
Construction of random forest (RF) and support vector machine (SVM) models. (A) Residual BoxPlot of RF and SVM, RF model showed a lower residual value; (B) Residual back-cumulative distribution diagram of RF and SVM, RF model showed a lower residual value; (C) ROC curve. The AUC of SVM is 0.857, The AUC of RF is 1.000; (D) A 10-fold cross-validation curve. Treat groups (red line), control groups (green line), and overall samples (black line). (E) The seven important CRGs.

### 3.4 Construction of the nomogram

We established a prediction model based on 3 significant PRGs via the “rms” package in R software (Figure 4A). The calibration curves in the results show the satisfactory accuracy of the new prediction model (Figure 4B). As the decision of DCA, figure 4C displayed the significant bias among the red, gray, as well as black lines associated with pyroptosis, which indicated the model’s prominent quality of decision-making capacity. The result of the clinical impact curve supported that the nomogram model constructed via PRG genes was of high precision and could efficiently predict the occurrence of OA (Figure 4D), which showed that the nomogram model was of considerable accuracy as well as reliability in predicting disease occurrence.

**FIGURE 4.**
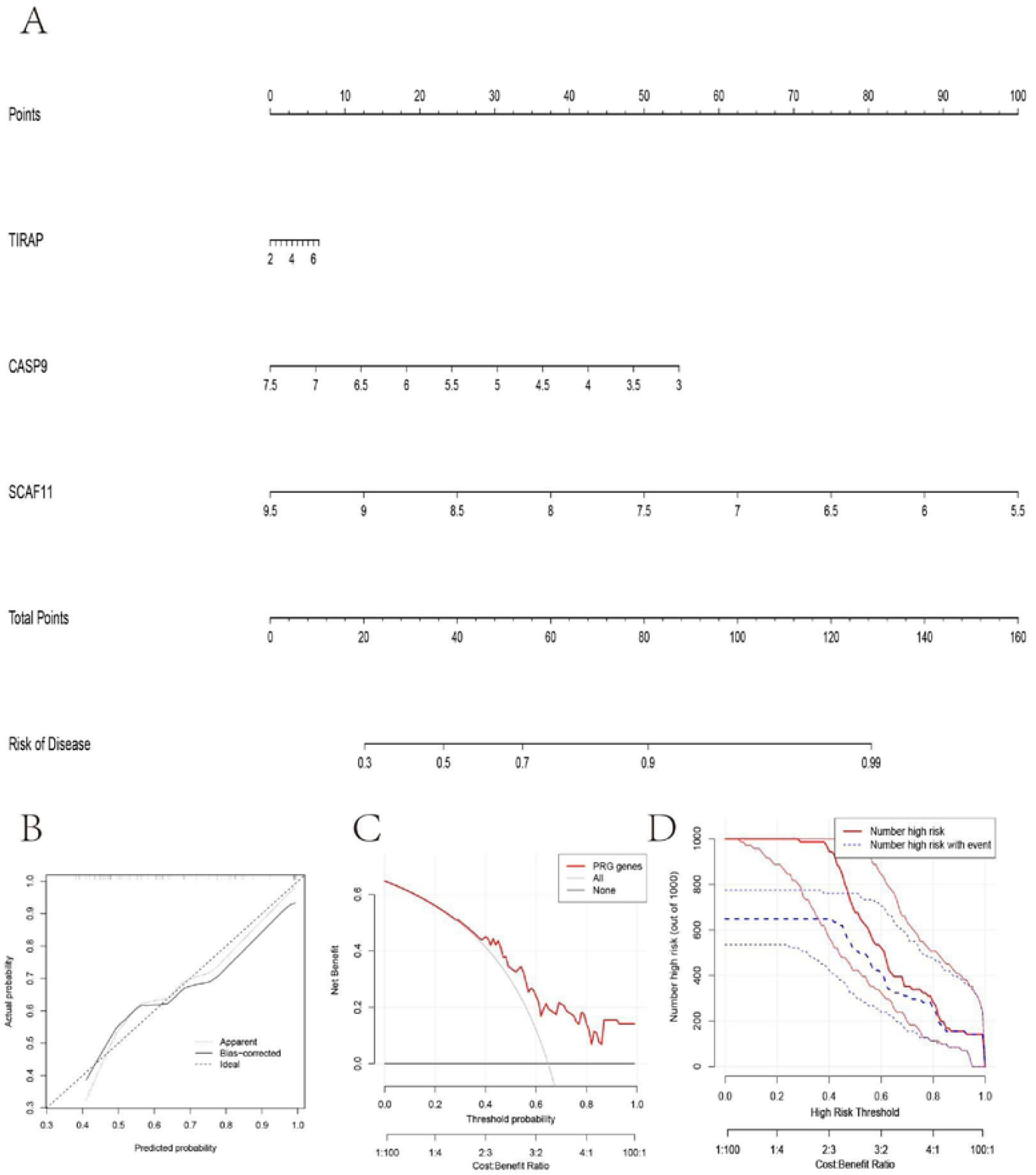
The nomogram model construction. (A) The nomogram of the model; (B) The calibration curve proves the accuracy of the prediction ability of the new model. (C) The red, gray, and black lines related to pyroptosis in the Decision Curve Analysis (DCA) are biased. (D) The clinical impact curve proves that the nomogram model can effectively predict the results with high precision.

### 3.5 Two PRG patterns identified by significant PRGs

According to the above significant PRGs, consensus clustering was conducted via the “ConsensusClusterPlus” package in R software. The separate pyroptosis patterns as well as two PRG patterns were confirmed (Figures 5A-C). Furthermore, the results showed that 7 PRGs, including CASP3, GPX4, NOD1, TIRAP, SCAF11, CASP4, and CASP9 in cluster B were of obviously higher expression levels than in cluster B (Figures 5D, E). Based on the results of PCA (Figure 5F), 7 PRGs could be grouped into two categories of PRG patterns.

**FIGURE 5.**
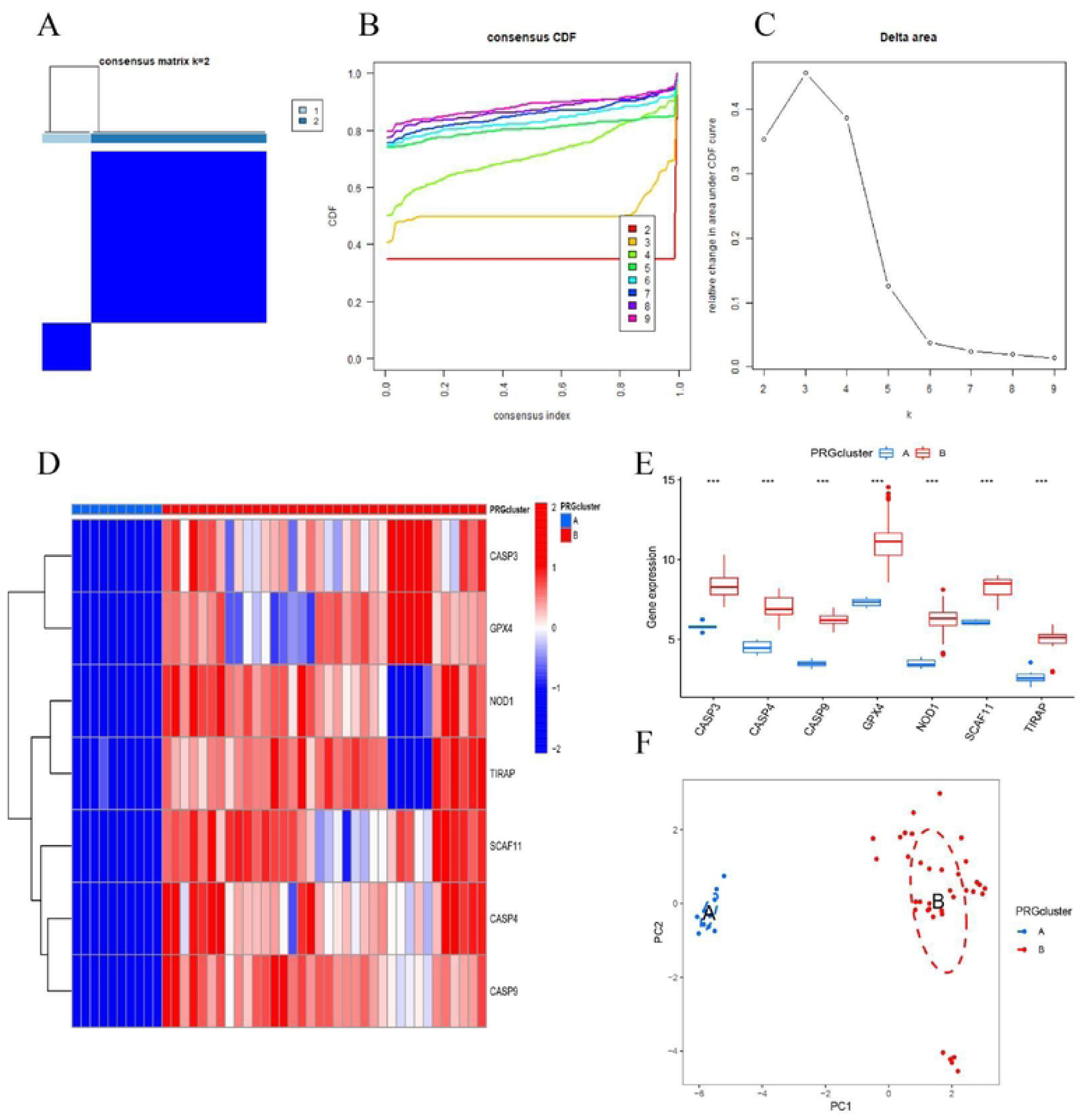
Consensus clustering of 7 PRGs in OA patients. (A) Consensus matrix of 7 PRGs with k = 2; (B) Cumulative distribution function (CDF) of consensus clustering; (C) Delta area plot of consensus clustering; (D) Differential expression heat map of 7 PRGs in cluster A and cluster B; (E) Expression histograms of 7 PRGs in cluster A and cluster B; (F) Principal component analysis (PCA) of 7 PRGs showed that the above genes could be divided into two PRG patterns. *P<0.05; **P<0.01; ***P<0.001.

Furtherly, the enrichment of immune cells of OA samples was explored via ssGSEA. Besides, the association of seven significant PRGs and immune cells was accessed, and the result was displayed in figures 6A–B, which indicated a positive association between GPX4 and abundant immune cells. We divided samples according to the GPX4 gene expression levels. High-expression of GPX4 samples were separated from the low-expression of GPX4 groups and subsequently explored the distinct immune cell infiltration patterns. Our results displayed that the immune cell infiltration increased in the subgroup with a high expression of GPX4 (Figure 6C). The results showed that cluster A may be associated with the OA’s incidence and development.

**FIGURE 6.**
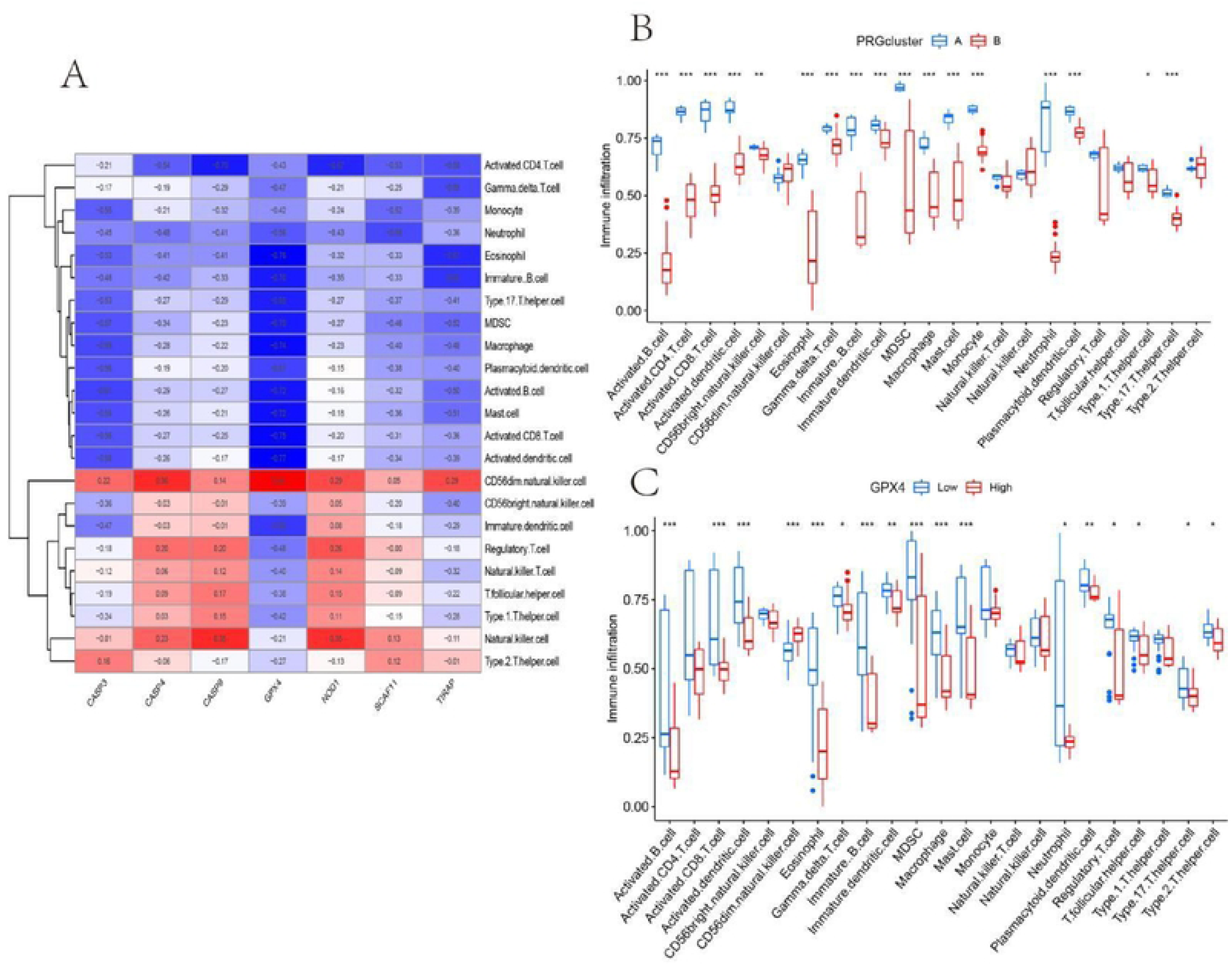
Analysis of immune cell infiltration. (A) The correlation between infiltrating immune cells and 7 PRGs; (B) Differential immune cell infiltration between cluster A and cluster B; (C) Comparison of immune cell abundance between high and low GPX4 expression group. *P<0.05; **P<0.01; ***P<0.001.

### 3.6 Identification of two divergent PRG patterns and PRG gene patterns

DEGs between clusters A and B were identified via the “limma” package in R software based on two significantly different PRG patterns above. Our results indicated that the genes were primarily enriched in cellular component sections, including cell leading edge (GO item ID:0031252), focal adhesion (GO item ID:0005925), the cell-substrate junction (GO item ID:0030055), collagen-containing extracellular matrix (GO item ID:0062023), and lamellipodium (GO item ID:0030027), and in molecular function sections including extracellular matrix structural constituent (GO item ID:0005201), growth factor binding (GO item ID:0019838), and integrin binding (GO item ID:0005178), and in biological process sections including cell-substrate adhesion (GO item ID:0031589), Golgi vesicle transport (GO item ID:0048193), regulation of cell morphogenesis (GO item ID:0022604), and regulation of cell-substrate adhesion (GO item ID:0010810) (Figures 7AC). Besides, the results of the KEGG pathway indicated that most DEGs were enriched in Human papillomavirus infection, Focal adhesion, Proteoglycans in cancer, Salmonella infection, Axon guidance, and the Hippo signaling pathway (Figures 7B).

**FIGURE 7.**
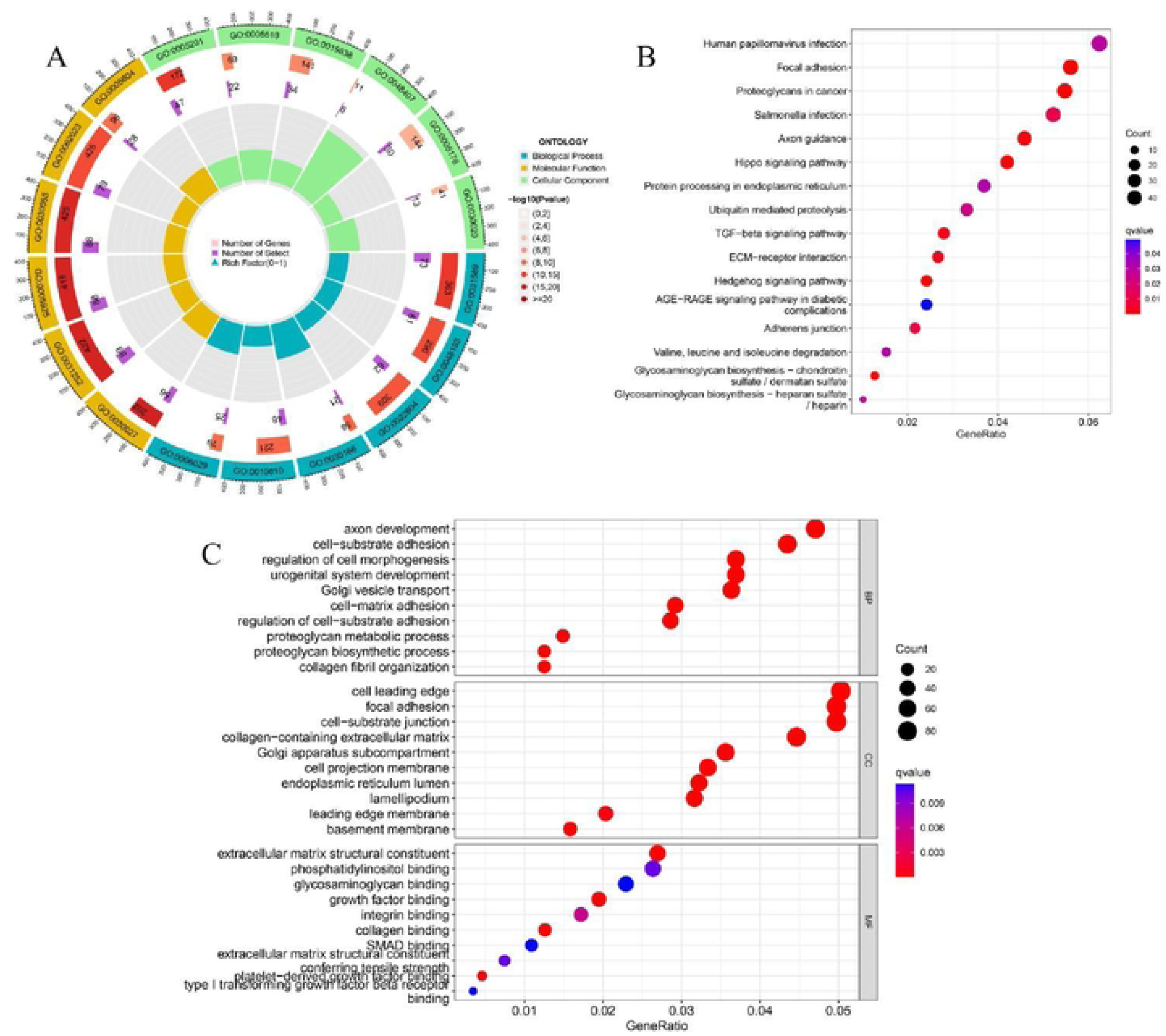
Enrichment analysis of DEGs between clusters (A, B), (A) GO circle diagram; (B) KEGG bubble chart; (C) GO bubble chart.

To further explore the PRG pattern, we organized OA patients into two different gene pattern subgroups by consensus clustering. Results indicated that different gene pattern subgroups were in conformity with the subgroups of PRG patterns (Figures 8A–C). Meanwhile, the expression level results of candidate PRGs and immune cell infiltration of the two gene clusters were shown in the boxplot, which was in line with the expression results of the previous PRG patterns (Figures 8D–E). Subsequently, to evaluate and compare two different PRG clusters’ PRG scores, the PRG pattern was quantified via the PCA algorithm. Results indicated that PRG cluster A is of higher risk score (Figure 9A). The results of the Sankey diagram (Figure 9B) displayed the association between PRG patterns and PRG scores. Also, the correlation between PRG patterns and OA was further explored to identify further the association between PRG patterns and cytokines (Figure 9C). We found that IL-4, IL-5, IL-13, TSLP, as well as IL-33 in the PRG cluster B were of higher expression levels.

**FIGURE 8.**
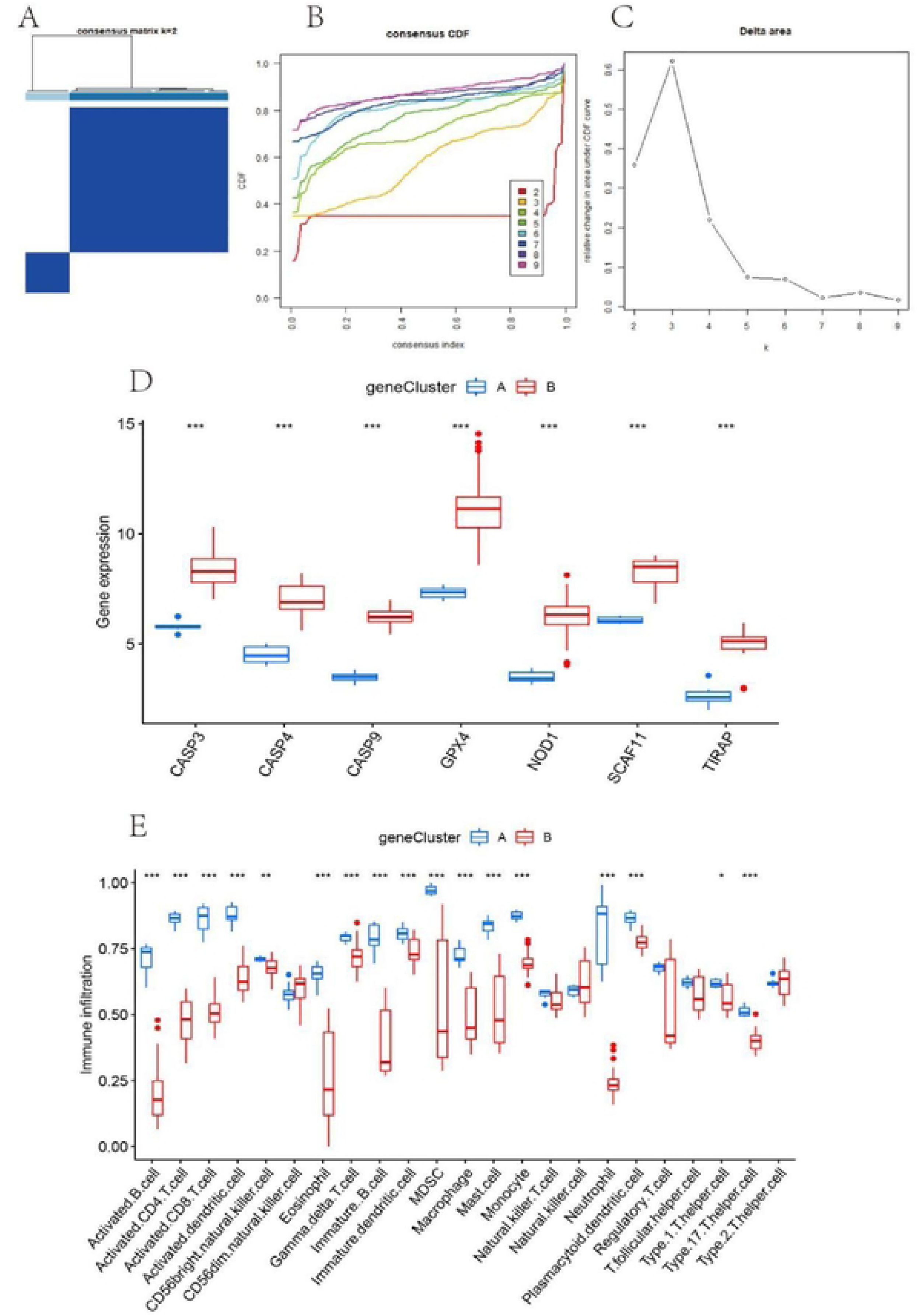
Consensus clustering of pyroptosis-related DEGs. (A) Consensus matrix of DEGs with k = 2; (B) CDF of consensus clustering; (C) Delta area plot of consensus clustering; (D) Expression histogram of 7 significant PRGs in cluster A and cluster B PRG gene models; (E) Different immune cell infiltration of DEGs in cluster A and cluster B PRG gene models. *P<0.05; **P<0.01; ***P<0.001

**FIGURE 9.**
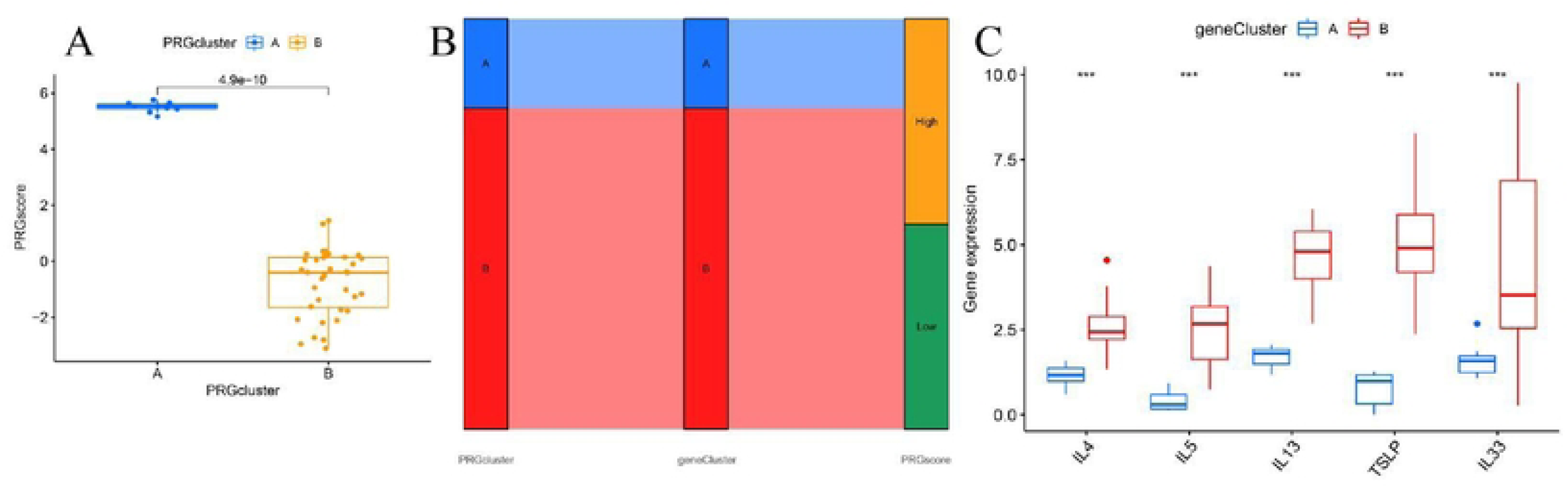
The role of different PRG patterns in diagnosing OA. (A) Differences in PRG scores between cluster A and cluster B; (C) Sankey diagram showing the relationship between PRG patterns; (D) Differential expression levels of interleukin (IL)-4, IL-5, IL-13, IL-33 and thymic stromal lymphopoietin (TSLP) between cluster A and cluster B. *P<0.05; **P<0.01; ***P<0.001.

### 3.7 qRT-PCR and immunohistochemistry

In order to verify the expression levels of the above key genes in OA cells and human cartilage tissues, qRT-PCR and immunohistochemical methods were used to detect the expression levels of CASP9, SCAF11 and TIRAP (primer sequences are shown in Table 1). Compared with normal rats, the expressions of CASP9, SCAF11 and TIRAP in LPS-treated rats chondrocytes were significantly decreased (P < 0.05), which was consistent with the above analysis results (Figure 10 A-C).Besides, 3 normal and 3 osteoarthritis cartilage samples were collected from the First Affiliated Hospital of Guangxi Medical University. Then, we detected and contrasted the expression of CASP9, SCAF11 and TIRAP by histochemical staining in two groups.And the immunohistochemical results also suggested similar results (Figure 11A-L).

**FIGURE 10.**
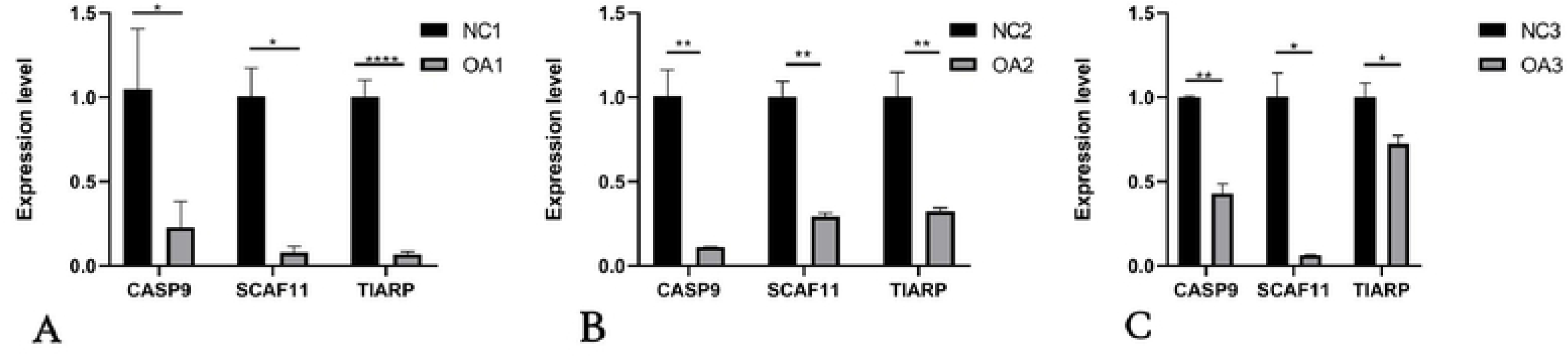
qRT-PCR analysis of three key genes in normal and osteoarthritis SD rat chondrocytes. (A-C) The expression of CASP9, SCAF11 and TIRAP in the cartilage of normal SD rats and OA rats. *P<0.05; **P<0.01; ***P<0.001.

**FIGURE 11.**
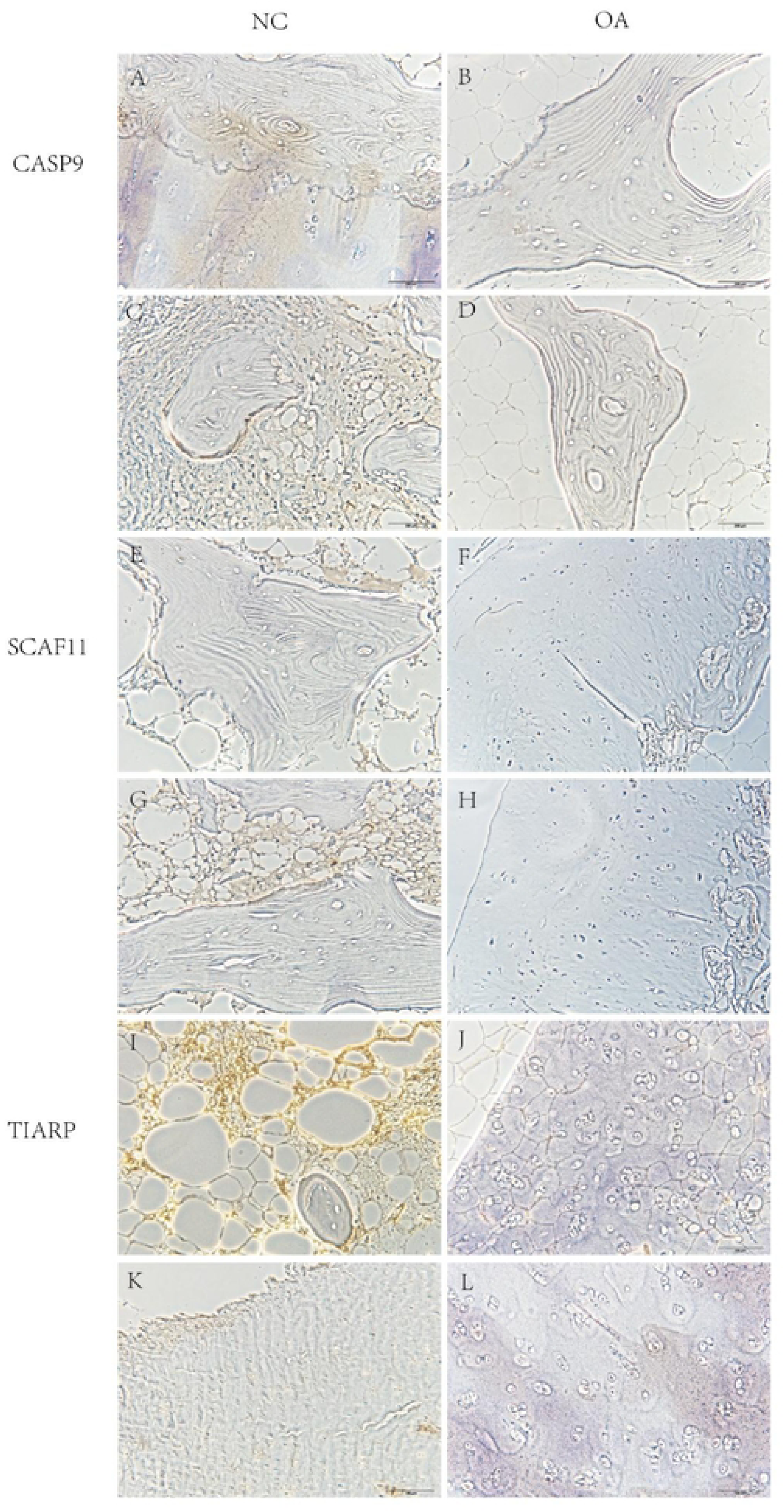
Immunohistochemical study of cartilage in normal people and patients with osteoarthritis.

**TABLE 1.**
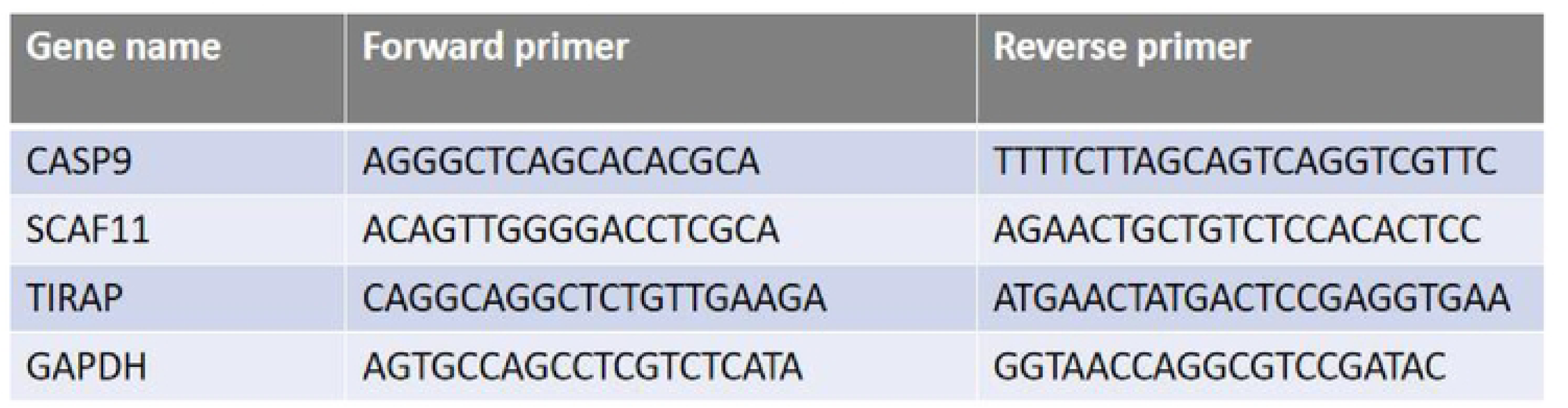
Primer sequences used in qRT-PCR experiments.

## 4 Discussion

We screened seven PRGs (CASP3, GPX4, NOD1, TIRAP, SCAF11, CASP4, and CASP9) in OA, and selected three characteristics of OA (TIRAP, SCAF11, CASP9) through random forest books, and constructed a nomogram of OA. In addition, we also verified the expression levels of these three characteristic genes by PCR and immunohistochemistry. In addition, we found two different burning patterns. We comprehensively evaluated the clinical significance of different models of pyroptosis. To further explore the infiltration level of immune cells and reveal the relationship between PRG and immune cell.

In terms of biochemical characteristics, the prominent markers of pyroptosis include the elements of inflammatory bodies, the activation of related caspases and gasdermin, as well as the release of a large number of pro-inflammatory elements, such as interleukin-1β (IL-1β), interleukin-18 and heat shock protein (9). In apoptotic cells, relevant caspases are activated to produce apoptotic bodies and factors such as Cyt c. Among them, the mediators of pyroptosis caspases involved factor caspase-1, caspase-4, caspase-5 as well as caspase-11, which have both initiator and effector functions. However, caspases involved in factor caspase-2, caspase-8, caspase-9 as well as caspase-10 initiate apoptosis and factor caspase-3, caspase-6 as well as caspase-7 perform apoptosis (10). It can be seen that the mechanisms of pyroptosis and apoptosis are different. Studies on pyroptosis and osteoarthritis have shown an important causal relationship.

We conducted the functional enrichment analysis of cluster A and subsequently via performing enrichment analysis employing DEGs between clusters A and B. The consequence of KEGG analysis indicated DEGs were obviously enriched in focal adhesion as well as the Hippo signaling pathway. The previous study has shown that the Hippo signaling pathway is composed of a series of conserved kinases, which is a signaling pathway that inhibits cell growth (11). In the metabolic process of the Hippo signaling pathway, the membrane protein receptor upstream was activated by the growth inhibition elements of the extracellular environment, goes through a group of kinase phosphorylation activation, and eventually plays on the downstream effector factors YAP as well as TAZ (12, 13). Hippo-YAP signaling pathway is a conserved and complex signal transduction pathway. Also, the YAP protein plays an essential role in cell signal transduction as well as connecting upstream and downstream factors(14). YAP protein is a pivotal transcription factor regulating cell fate in the Hippo signaling pathway, and studies have shown that YAP protein promotes cell proliferation and differentiation (15). Zhong et al. (16) found that down-regulating the expression of YAP could help maintain the phenotype of chondrocytes and inhibit the proliferation of chondrocytes. Karystinou et al. (17) showed that YAP can directly regulate BMP and transform growth factor signals through the interaction of Smad protein. YAP may be a negative regulator of chondrocytes, which plays a role by inhibiting chondrocyte signal transduction and activating proliferation activity simultaneously. Hippo-YAP signaling pathway and chondrocyte apoptosis: Hippo-YAP signaling pathway affects apoptosis by receiving the membrane FAT4 membrane protein receptor signal and transmitting it to the YAP protein through various membrane proteins. YAP protein transduces the anti-apoptotic signal to the nucleus. Finally, it regulates the effects of anti-apoptotic factors BCL-2, factor AKT and factor caspase-3, indicating that YAP protein may be an important factor promoting the apoptosis of articular cartilage during the onset of osteoarthritis.

The programmed inflammatory cell death, pyroptosis, may play an important role when chondrocyte apoptosis cannot be properly activated, and cell death does occur. Unlike apoptosis and simple cell necrosis, pyroptosis is pro-inflammatory (18). Studies have shown that there may be chondrocyte pyroptosis-mediated chondrogenic inflammation and matrix degradation in OA chondrocytes (19). In addition, there is growing evidence that pyroptosis may occur in various tissues associated with chronic aseptic inflammation and tissue fibrosis. For example, pyroptosis of microglia can lead to neurogenic inflammation and fibrosis (20), and pyroptosis of renal macrophages can lead to renal inflammation and fibrosis (21). As inflammatory factors, such as interleukin-1β (IL-1β) as well as IL-18 have been recognized to enhance cartilage catabolism and promote the degradation of chondrocyte extracellular matrix by inhibiting the production of proteoglycan and type II collagen, thus leading to the occurrence as well as progression of OA (22). The previous studies have indicated that pro-inflammatory elements such as IL-1β as well as IL-18 secreted by scorch death of synovial macrophages can directly cause synovial inflammation and participate in inducing cartilage matrix degradation, promoting the pathological process of OA and inhibiting scorch death of synovial macrophages can alleviate synovitis and fibrosis in OA (21).

Our research identified two PRG clusters. Results indicated that immune infiltration was much more obvious in cluster A, manifesting that patients in cluster A may have more obvious benefits in the treatment of OA disease. Our research found that macrophages, dendritic cells (DC), and T cells significantly differed between the two groups.

Pattern recognition receptors (PRRS) on the surface of macrophages recognize PAMP and DAMP in the joint lumen, bind to TLR/IL-1 receptors and activate the NF-κB pathway to up-regulate the release of inflammatory factors. Another important activation pathway is through the inflammasome produced by cells under stress, which binds to nucleotide-binding oligomerized domain-like receptors, thereby releasing inflammatory factors. Macrophages can also be activated and differentiated into M2 subgroups through selective pathways, thus showing the role of inhibiting inflammation and promoting tissue repair (23). Activated macrophages can highly express mannose-receptor (CD206) under the stimulation of IL-4 and IL-13, as well as IL-4 can stimulate the transformation of arginine to ornithine in macrophages, thereby increasing the secretion of cell-matrix components such as collagen as well as polyamine proteoglycans (24). The weakened repair function of the M2 subgroup can lead to the persistence of tissue damage, and the strong repair function can lead to tissue fibrosis (25). Activated macrophages can highly express folate receptors. Piscaer et al. (26) used SPECT/CT to trace ^111^INCL3-labeled folate receptors to observe the rat OA model prepared by iodoacetate injection and anterior cross-ligament amputation, respectively. The results showed strong folate receptor signal aggregation in the knee joint of rats, suggesting the important role of activated macrophages in OA.

OA is portrayed as intra-articular immune cell infiltration (27). Existing evidence shows that immune cells in OA synovial tissue (especially in the synovial tissue with obvious inflammation) mainly include macrophages, T cells, B cells, natural killer cells, as well as dendritic cells, etc. (28). Among them, macrophages accounted for 65% and T cells accounted for 22% (29). Symons et al. (30)detected soluble CD4 in OA joint fluid by ELISA. They found that the concentration of soluble CD4 in OA patients was lower than that in rheumatoid arthritis patients but higher than that in normal people. Hussein et al. (31) compared the proportions of CD4+ as well as CD8+ T cells in the synovial fluid of OA and rheumatoid arthritis patients by flow cytometry. They found that the proportions of CD4+/CD8+ were similar, and CD4+ T cells were dominant. Kriegova et al. (32) also obtained the same result and found that female patients with CD4+ T cells were taller. These results indicate that T cells were tightly associated with the pathological process of OA. The mechanism of T cells participating in the pathological process of OA may be related to its recognition of the breakdown products of cartilage matrix proteoglycans. de Jong et al. (33) found that T cells can specifically recognize proteoglycan 16-39 and 263-282 amino acid sites, thereby increasing the release of inflammatory factors. Current research evidence suggests that it is related to OA pathology. The related cell subsets were mainly Th1, Th9 and Th17 cells. Studies have compared the proportion of Th1/Th17 cells in circumference blood, joint fluid as well as synovium of OA patients with rheumatoid arthritis and found that the proportion of Th1/Th17 cells in circumference blood and synovium of OA patients is higher, with more Th1 cells in synovium and fewer Th17 cells (34, 35). As a powerful APC, DC is closely related to the destruction and repair of articular cartilage. Although the distribution of DC in human joint fluid and bone marrow is small, its content is positively correlated with the severity of OA. Matrix metalloproteinase (MMP), as the main collagenase component of OA cartilage, is closely related to the number of plasmacytoid DC (p DC) subpopulations of DC. DC can facilitate the differentiation of B-type synovial cells as well as plays an essential part in the OA’s early stages.

The main shortcomings of this article are as follows. Firstly, no data from in vivo, in vitro research, or clinical studies could be used to corroborate the result. What’s more, the level of proteomics and metabolomics is not reflected in the evaluation of pyroptosis based on mRNA level alone.

## Conclusion

This research constructed a diagnostic practical model according to PRGs in OA and fully explored the relationship between pyroptosis and immune function. Three characteristic genes were also validated in PCR and immunohistochemistry. In actual practice, the predictive significance of pyroptosis death risk score during therapy in OA patients deserves further study.

## Data availability statement

The original contributions presented in the study are included in the article/supplementary material. Further inquiries can be directed to the corresponding author.

## Author contribution

Jun Yao designed the study and revised the manuscript. Meimei Xiao, Yue Qiu and Jinzhi Meng conducted data analysis and experiments. Cancai Jiang and Yanming Chen wrote a draft manuscript. All authors contributed to the article and approved the submitted version.

## Acknowledgments

The authors thank the authors who provided the GEO public datasets.

## Publisher’s note

All claims expressed in this article are solely those of the authors and do not necessarily represent those of their affiliated organizations, or those of the publisher, the editors and the reviewers. Any product that may be evaluated in this article, or claim its manufacturer may make, is not guaranteed or endorsed by the publisher.

